# Single-cell interactomes of the human brain reveal cell-type specific convergence of brain disorders

**DOI:** 10.1101/586859

**Authors:** Shahin Mohammadi, Jose Davila-Velderrain, Manolis Kellis

## Abstract

The reference human interactome has been instrumental in the systems-level study of the molecular inner workings of the cell, providing a framework to analyze the network context of disease associated gene perturbations. However, reference organismal interactomes do not capture the tissue- and cell type-specific context in which proteins and modules preferentially act. Emerging single-cell profiling technologies, which survey the transcriptional cell-state distribution of complex tissues, could be used to infer the single-cell context of gene interactions. Here we introduce SCINET (Single-Cell Imputation and NETwork construction), a computational framework that reconstructs an ensemble of cell type-specific interactomes by integrating a global, context-independent reference interactome with a single-cell gene expression profile. SCINET addresses technical challenges of single-cell data by robustly imputing, transforming, and normalizing the initially noisy and sparse expression data. Subsequently, cell-level gene interaction probabilities and group-level gene interaction strengths are computed, resulting in cell type specific interactomes. We use SCINET to analyze the human cortex, reconstructing interactomes for the major cell types of the adult human brain. We identify network neighborhoods composed of topologically-specific genes that are central for cell-type influence but not for global interactome connectivity. We use the reconstructed interactomes to analyze the specificity and modularity of perturbations associated with neurodegenerative, neuropsychiatric, and neoplastic brain disorders; finding high variability across diseases, yet overall consistency in patterns of cell-type convergence for diseases of the same group. We infer for each disorder group disease gene networks with preferential cell-type specific activity that can aid the design and interpretation of cell-type resolution experiments. Finally, focusing on the pleiotropy of schizophrenia and bipolar disorder, we show how cell type specific interactomes enable the identification of disease genes with preferential influence on neuronal, glial, or glial-neuronal cells. The SCINET framework is applicable to any organism, cell-type/tissue, and reference network; it is freely available at https://github.com/shmohammadi86/SCINET.

## Introduction

Proteins participate in crosstalking pathways and overlapping functional modules that collectively mediate cell behavior. The complete set of molecular components and interactions form an “interactome,” which, although incompletely characterized, has provided a reference structure for the systems-level study of multiple organisms, and a core framework for network biology (Barabási and Oltvai 2004). In the past decade, large efforts have reconstructed multiple organismal reference networks (Li et al. 2017; Rolland et al. 2014). The study of the human interactome has been instrumental in revealing the structural context, modularity, and potential mechanisms of action of disease associated perturbations (Loscalzo 2017; Menche et al. 2015), which often tend to disrupt protein-protein interactions (Sahni et al. 2015).

Reference organismal networks, however, do not provide information about the specific spatiotemporal context in which gene interactions might occur, prohibiting the direct study of the context where the effect of their perturbation most likely manifests. One approach to incorporate context into a global interactome network is by considering the transcriptional dynamics of the network’s components, effectively integrating a static snapshot of the space of all potential interactions with context-specific gene expression. This approach has been previously applied to construct tissue-specific networks, taking advantage of bulk gene expression measurements (Mohammadi and Grama 2016; Magger et al. 2012; Bossi and Lehner 2009). With the recent development and increasing use of single-cell technologies, it is now possible to profile large-scale cell atlases of heterogeneous tissues and cell-types across multiple organisms (Tabula Muris Consortium et al. 2018; Regev et al. 2017; Rozenblatt-Rosen et al. 2017). The increasing availability of these data provide a unique opportunity to study the context-specificity of molecular interactions at single-cell resolution. A naive approach to infer cell type-specific networks would be to adopt techniques developed for bulk expression data. In practice, however, single-cell datasets are extremely sparse, with many genes having zero expression values due to both biological and technical factors (Angerer et al. 2017). Moreover, single-cell profiles, for the first time, enable estimatimating a distribution of interaction strengths within a cell group, directly assessing inter-cellular interaction variability. Finally, the increasing resolution comes at a computational cost, due to the increasing size of cell-level data. Thus, the development of highly efficient yet accurate techniques is required to transition to a single-cell network biology.

Here we introduce SCINET (Single-Cell Imputation and NETwork construction), a computational framework that overcomes technical limitations in single-cell data analysis, enabling the reconstruction of cell- and cell-type specific interactomes by integrating a reference interactome with single-cell gene expression data (Figure 1a). SCINET includes multiple features that directly meet challenges associated with single-cell analysis. First, a regression-based imputation step circumvents the high level of noise and sparsity intrinsic to single-cell data by inferring missing values and balancing gene expression levels. Second, a rank-based inverse normal transformation accounts for the drastic differences in the distribution of different genes, resulting on comparable expression scales. Third, a novel statistical framework with analytical closed-form solution enables efficient inference of gene interaction likelihoods. Finally, a subsampling scheme allows the computation of cell-type (group) level interaction strengths, despite cell-level data incompleteness. All together, SCINET simply takes as input a reference interactome network, a scRNA-Seq count matrix, and corresponding cell-type (group) annotations; and it returns a set of cell-type (group) specific networks, in which every interaction is represented by its estimated mean and standard deviation across the samples of each group.

**Figure 1.**
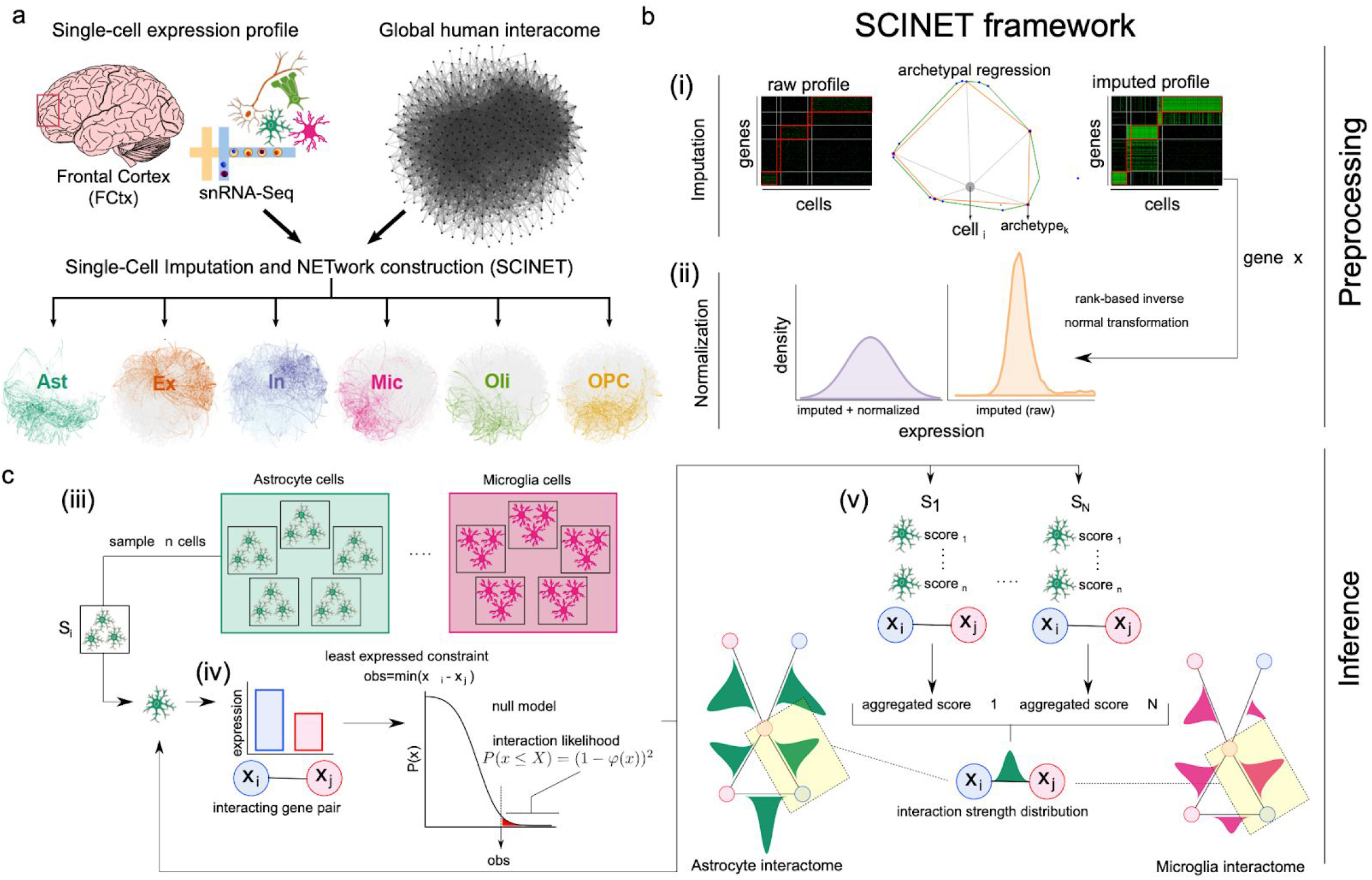
Overview of SCINET. **a**, SCINET integrates a reference interactome network with a cell-annotated single-cell transcriptomic dataset to reconstruct cell-group specific interactomes. In this study we applied SCINET to produce cell-type interactomes for the major cell-types of the adult human frontal cortex. **b**, SCINET preprocesses single-cell expression by first using a matrix decomposition method to interpolate values for missing observations. Subsequently, the distribution of expression values is transformed into a common and comparable distribution using rank-based inverse normal transformation. Preprocessing results in across-gene comparable gene activity scores. **c**, SCINET infers interaction strength distributions for each interaction and cell-type(group) using a novel statistical framework. A sample of n cells of a certain type is randomly chosen from the cell-type population. For each cell, the minimum activity score of each pair of interacting genes is used to quantify the strength of a potential interaction based on the tail of an analytical null model measuring the likelihood of observing such value under independence. The scores computed for the n samples cells are then aggregated into one aggregate score (meta p-value). The sampling procedure is repeated a large number of times. Finally, the mean and variance of the distribution of aggregate scores is recorded for each gene interaction and cell-type (group).

Using recently-published single-nucleus RNA-Seq (snRNA-Seq) data from 10,319 cells of the human prefrontal cortex (Lake et al. 2018), and an integrative reference interactome combining data from 21 gene interaction resources (Huang et al. 2018), we applied SCINET to infer networks specific to six major cell-types of the human prefrontal cortex; resulting in astrocyte, excitatory neuron, inhibitory neuron, microglia, oligodendrocyte, and oligodendrocyte progenitor interactome networks. These networks enable the distinction of *topologically-specific genes,* whose overall interaction strength is highly cell-type-specific, vs. *topologically-invariant* genes, whose connectivity pattern is not significantly influenced by the cell-type context. We found that *topologically-specific* genes exhibit cell-type related functions and overlap with known markers, suggesting that inferred topological properties recover the contextual influence of otherwise noncentral genes.

We then used the set of cell-type specific networks to study the network context of 19,250 disease-gene associations, including 29 diseases and 6,979 genes. We evaluated whether perturbations associated with brain-related disorders show significant cell-type specific modularity, measured using a novel random-walk-based approach that assessed the strength of the local connectivity of disease genes in each network, relative to same-sized random gene-sets. We found a strong but variable modularity among disease genes across different disorders. Moreover, we found that individual diseases show variable modularity across cell-types, a property that we used to define cell-type perturbation profiles for the disorders. The computed profiles showed overall high similarity across phenotypically-related disorders, with strong preferential modularity in cell-types consistent with the biology of the disease. We found that neurodegenerative disorders preferentially target astrocyte and microglia cells, while psychiatric disorders strongly target neuronal specific interactions. Consistent with biological expectation, neoplastic disorders affect interactions occurring on cells with proliferative potential (i.e., astrocyte and oligodendrocyte progenitor cells).

Finally, we investigated whether cell-type specific gene prioritization could discriminate shared and cell-type specific components of disease-associated perturbations within the network context. Using schizophrenia and bipolar disorder as a case study, we found that genes converge to modular network regions enriched with disease relevant but distinct processes, exemplifying a systems view of the pleiotropic effect of gene sets associated with complex disorders.

Overall, our approach provides a general framework to study the context-specificity of global interactome networks using single-cell transcriptional profiles. It can be used to integrate and study any combination of reference network(s) and single-cell activity profiles.

## Results

### Methodological overview

The core SCINET framework (Figure 1) is based on the following novel developments: (i) a decomposition method to interpolate values for missing observations in the scRNA-Seq profile, (ii) a parametric approach to project heterogeneous gene expression distributions into a compatible subspace (Figure 1b), (iii) a statistical framework to measure the likelihood of gene interactions within each cell, and (iv) a subsampling approach to aggregate interaction likelihoods of individual cells, reduce noise, and to estimate the underlying distribution and variability of interaction strengths within each cell-type population (Figure 1c).

The first component (i) is built upon our previously developed method ACTION (Mohammadi et al. 2018). Briefly, a single-cell expression matrix is iteratively decomposed into lower dimensional matrices at different levels of granularity defined by a number of low-rank factors *(archetypes).* This results in a small set of landmark transcriptional states (patterns) that optimally represent the variability in the dataset. We then use the set of discovered patterns to interpolate transcriptional profiles for individual cells (Figure 1b, i) (for details, see Methods). The second component (ii) was designed to overcome the fundamental differences in the expression distribution of genes, and their particular skewed distribution across single cells. The original values do not allow a direct comparison of the distributions of different genes. We approached the problem by using a rank-based inverse normal transformation to rescale the gene expression distributions and project them into a common, normally distributed subspace. We refer to the resulting interpolated and normalized expression profiles as gene activity scores (Figure 1b, ii). Components (iii) and (iv) introduce a statistical framework that maps gene activity scores to the reference interactome. We assume that the feasibility of occurrence of an interaction within a cell is dictated by the interacting partner with the least activity. To formalize this notion, we use the minimum activity score of each pair of interacting genes as a statistic to assess the strength potential of the interaction in a given cell. We quantify such potential for each interaction pair and cell using the tail of an analytical null model measuring the likelihood of observing a certain interaction strength under independence (see Methods). We then aggregate strength scores by combining the individual likelihood values within random subsamples of cells into a meta *p*-value using Fisher’s method (Fisher 2006). Finally, to estimate the variability of the interaction strength scores, and to account for the inherent noise of single-cell activity, component (iv) implements an ensemble learning technique based on random subsampling that estimates an interaction strength distribution for each interaction and cell-type (Figure 1b, iii-v).

### Constructing interactomes specific to the major cell-types of the human cortex

To demonstrate the applicability of SCINET, we used single-cell expression data of the human prefrontal cortex reported in (Lake et al. 2018) to infer cell-type specific interactomes. After removing endothelial cells and pericytes (due to small size) and independently combining cells annotated as subtypes of excitatory or inhibitory neurons, we defined a set of cell-type annotations considering the 6 major cell-types of the brain: astrocytes, excitatory neurons, inhibitory neurons, microglia, oligodendrocytes, and oligodendrocyte progenitor cells (OPCs). The dataset includes the expression values of 22,002 genes across 10,223 cells. Among these genes, 6,822 were expressed in less than ten cells and were removed from the study, resulting in a total of 15,180 retained genes. We selected as reference human interactome a recently curated gene network, the *parsimonious composite network (PCNet) (Huang et al. 2018).* PCNet was constructed by integrating 21 network databases, retaining only interactions supported by at least two independent sources; it includes 2,724,724 (2.7M) interactions among 19,781 genes. We intersected PCNet with the filtered expression profile, ending up with a network of 1,882,141 (1.8M) interactions among 13,581 genes, which we used for the remainder of the study.

We applied SCINET to the matched scRNA-Seq count matrix and PCNet network to reconstruct 6 cell-type specific networks. These networks have between 4886 and 7331 genes (35.4% and 54.6% of the original PCNet network), with the smallest number of retained genes observed in the oligodendrocyte network and the largest in the excitatory neuron network. For this reconstruction, we considered a bonferroni-adjusted *p*-value threshold of 0.01 to prune significant edges. After pruning, and retaining only the largest connected component of each network, we obtained a final set of networks including between 4,416 (OPC) and 6,435 (Ex neurons) genes connected by a subset of interactions corresponding to 13.2-30.5% of total interactions in the reference PCNet (Figure. 2a). The obtained networks show topological properties similar to those of other complex biological networks, namely skewed degree distributions with a small number of highly connected genes (Supplementary Figure 1). The vast majority of gene interactions occur only in one cell-type (private interactions), indicating that SCINET provides high specificity. One exception is interactions shared between Ex and In neurons (neuronal interactions), which account for ~18% of the interactions observed across neurons. Similarly, Ast and OPC networks share ~15% of the total interactions observed in these two glial cell types (Supplementary Figure 2a). The two neuronal and two glial progenitor cells (Ast and OPC) respectively share functional properties, which are possibly captured by common interactions recovered by SCINET. Overall, SCINET recovers cell-type specific gene networks whose patterns of shared interactions capture similarities and differences among neuronal and glial populations (Supplementary Figure 2b). Importantly, we corroborated the robustness of the inferred networks to gene expression interpolation (SCINET step 1), observing significant overlap of inferred cell-type specific interactions across existing scRNA imputation methods (Supplementary Figure 3). SCINET brain cell-type specific interactomes are reported in Supplementary file 1.

**Figure 2.**
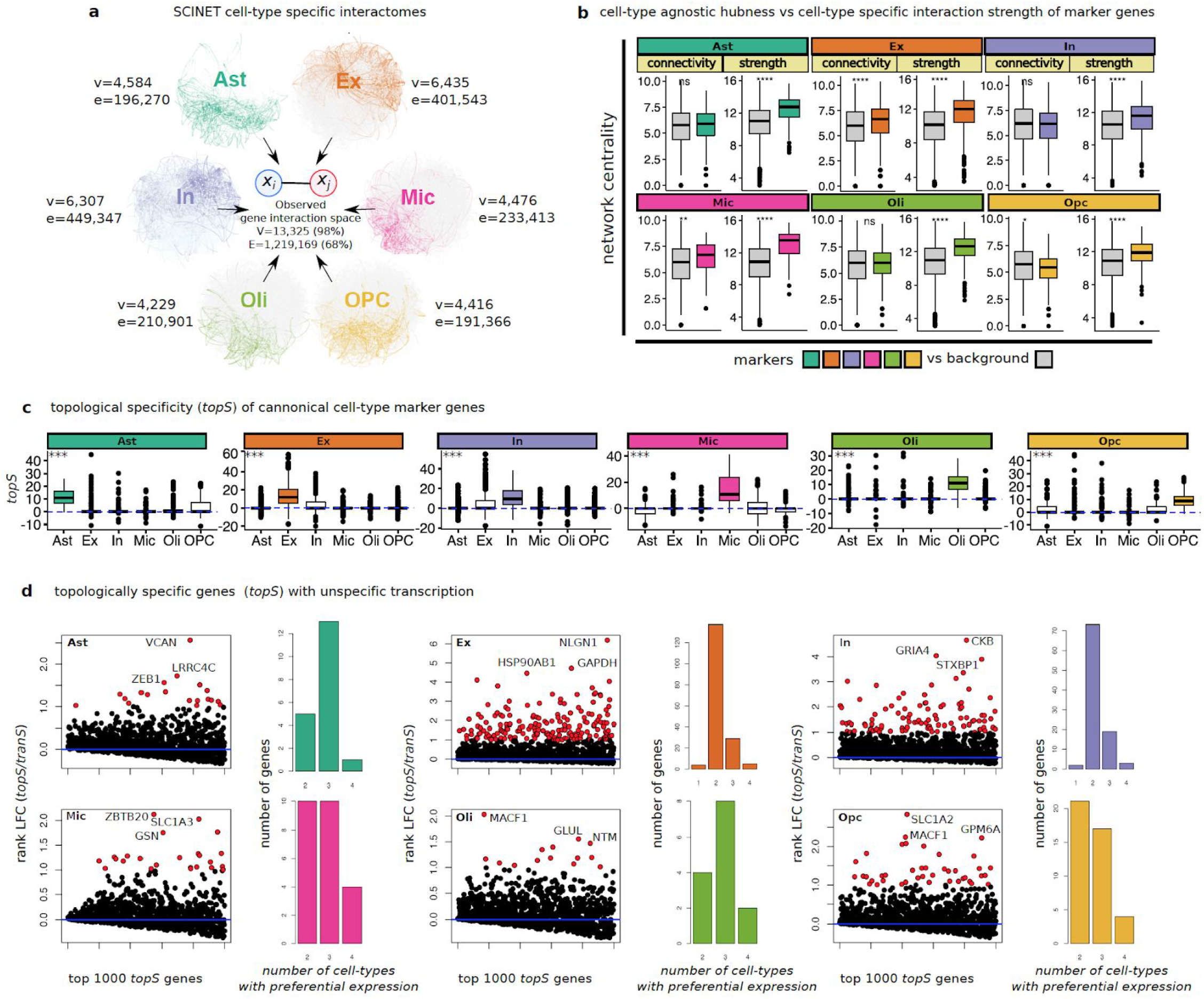
Patterns of specificity of identified interactions. **a**, summary of inferred cell-type specific networks. **b**, patterns of connectivity and interaction strength computed within cell-type specific networks for canonical markers of each cell-type. Statistical test: two-sided Wilcoxon rank sum test. **c**, distribution of topological-specificity scores computed for each cell-type specific network for canonical transcriptional markers of a particular cell-type. Statistical test: Kruskal–Wallis one-way analysis of variance. **d**, topologically specific genes that show promiscuous expression across cell-types. Convention for symbols indicating statistical significance: ns: p > 0.05, *: p <= 0.05, **: p <= 0.01, ***: p <= 0.001, ****: p <= 0.0001.

### Topological-specificity highlights genes with preferential cell-type influence

In network analysis, it is intuitively considered that highly connected nodes (i.e., hubs) are major determinants of systems behavior (Kitsak et al. 2010). In fact, hub proteins of reference interactomes in model organisms have been associated with properties suggestive of functional influence and constraint, such as essentiality, conservation, and phenotypic effects upon perturbation (Barabási and Oltvai 2004). Starting from a reference interactome, where “hubness” is defined based on the invariant number of total interactions for a node, SCINET infers cell-type specific networks; where dynamic interaction strengths capture cell-type specific transcription. We sought to study the relationship between cell-type specific hubness and the context-specific role of the corresponding genes.

First we examined whether known canonical cell-type markers have a distinctive topological role in the networks. We found that total connectivity alone does not discriminate markers from non-marker genes. In other words, with the exception of Ex neurons, markers do not show a significantly higher number of interactions in their corresponding cell-type, relative to non-marker genes. On the other hand, in all cases, markers do show a significantly higher aggregate interaction strength (sum of SCINET inferred interactions strengths) relative to non-markers in the same cell type (Figure 2b). This observation suggests that SCINET is able to infer context specific influence as measured by interaction strengths.

To directly quantify the influence of a gene in a given context, while decoupling the effect of its reference connectivity, we introduce a measure of gene *topological-specificity* (*topS*). *topS* decouples the two centrality measures (connectivity and strength) by measuring the deviation of the observed overall strength of the interactions of a gene in a given cell-type, relative to the expected strength under a random model that preserves the reference topology of the interactome but reshuffles cell-type specific interaction strengths (see methods). We tested whether curated cell-type marker genes tend to rank high in *topS* for the corresponding cell-type and not for the others. We found that marker genes, indeed, exhibit a significantly higher *topS* in the corresponding cell-type (Figure 2c), indicating that the inferred local interaction strengths capture cell-type relevant biological roles that are not explained by reference connectivity alone. Consistent with the cell-type specific statistical shift in *topS* scores, we found that many of the top ten *topS* genes for each cell-type are known canonical markers of the corresponding cell-type (Supplementary Table 1).

Marker genes by definition are transcriptionally specific -- i.e., are specifically expressed in the cell-type in question. It was previously shown that constitutive proteins may acquire context-specific effects by means of tissue-specific interactions (Bossi and Lehner 2009). We reasoned that *topS* will capture such counterintuitive property, even when globally both transcriptional and topological specificity correlate (Supplementary Figure 4). We used *topS* scores to systematically define genes with a preferentially influential role in a cell-type network that is not explained solely by their expression pattern. By comparing a gene’s measure of transcriptional specificity *(tranS)* with its *topS,* we identified groups of genes with high *topS* and low *tranS* (Figure 2d). These genes are expressed in multiple cell-types, but, through patterns of cell-type specific interactions, play differential centrality roles across the networks. Among the top 3 genes with the strongest deviation in *topS* relative to *tranS* per cell-type, we identify, for example, genes involved in generic functions such as protein folding (HSP90AB1) and energy homeostasis (CKB); as well as genes with more specific functions such as membrane transporters with activity in multiple glial cells (SLC1A2, SLC1A3). Transcriptional and topological specificity scores are reported in Supplementary table 2.

These results show examples of how the *topS* measure of SCINET can be used to measure for each gene a network influence score that is context-specific but not explained by either the gene’s reference connectivity or expression specificity alone. Instead, differential patterns of interactions determine the context specific influence of promiscuously expressed genes.

### Disease-associated genes tend to be connected by cell-type specific interactions

Next we used the inferred networks to study the patterns of gene interaction among disease associated genes. Previous studies have shown that disease associated genes tend to form coherent neighborhoods in the human interactome (Menche et al. 2015). That is, disease genes tend to connect to each other directly or through intermediate interaction partners. We ask whether genes associated with certain disease classes of brain disorders have system-level preferential influence on specific cell-types, as evidence by (1) preferential occurrence of interactions between disease genes in given cell-types, and (2) a more coherent modular connectivity among genes independently associated with similar disorders in a given cell-type.

We collected, curated, and annotated gene-disease associations for brain-related disorders by matching the DisGeNET database (Piñero et al. 2017) with annotations from the Monarch Disease Ontology (MONDO) (Bello et al. 2018). We organize diseases in three separate classes: (i) neurodegenerative, (ii) neuropsychiatric, and (iii) neoplastic disorders. In total, we analysed 5,069 protein coding genes that have been associated with at least one of 29 brain disorders: 5 neurodegenerative, 6 neoplastic, and 18 psychiatric (Supplementary Table 3). The genes analyzed are those present in both DisGeNET and PCNet.

Using these “disease genes”, we constructed for each disease a subnetwork by extracting from the global interactome only the disease-associated genes and the direct interactions among them. We then aggregated the set of interactions for all diseases within each class, resulting in 202,598, 203,324, and 263,950 interactions for neurodegenerative, neuropsychiatric, and neoplastic disorders, respectively. These disease class networks represent context-agnostic interactions among disease-associated genes supported by PCNet. To assess whether gene interactions for different disease classes preferentially occur in specific cell-types, we defined sets of cell-type specific interactions inferred by SCINET (adjusted p-value<0.01), that we used to evaluate their overrepresentation within interactions linking disease group genes. We considered independently each pair of disease class/cell type (Figure 3a). Overall, our results suggest that cell-type specific interactions are more likely to be perturbed in disease relative to nonspecific interactions, with preferential cell-types being targeted by the different disease groups. The first clear group distinction is that gene interactions associated with neurodegenerative and neoplastic disorders target glial cells, whereas neuropsychiatric disorder genes almost predominantly interact in neuronal cells (Figure 3a). In terms of specific cell-types, we found that neurodegenerative associated gene interactions are significantly enriched among cell-type interactions in microglia and astrocyte specific networks. Neoplastic disorders are depleted among neuron-specific interactions and overrepresented in astrocyte and OPC interactions. Neuropsychiatric disorders are strongly enriched in excitatory-neuron interactions. For each of the dominant cell-types by disease class, we extracted the top 10 strongest interactions, and found that these interactions in each case cluster in only one or two connected modules (Figure 3b). These modules do not share genes across cell-types; however, the association of the same cell-type with multiple disease types is mediated by different genes interacting in the same cell-type. We found Astrocytes as a cell-type strongly associated with both neurodegenerative and neoplastic disorders. Among the top astrocyte gene interactions, two interactions involving the gene CLU (CLU-GLUL and CLU-CST3) are associated with both disorder types. CLU is an example of a highly pleiotropic gene, which is associated with hundreds of diseases (according to DisGeNET). In the neurodegenerative context, CLU plays a chaperone role important for the prevention of amyloid aggregation and fibril formation. It also, however, plays a role in cell proliferation of relevance for neoplasia. This multiplicity of function might stem from differential use and perturbation of gene interactions, which could be more extensively studied using SCINET. In this example, we observed that a subset of CLU strong gene interactions occurring in astrocytes, distinguish associations with neoplasia (CPE, GJA1, and FGFR3) or with neurodegeneration (PON2, WWOX). Networks including gene interactions between disease class genes that overlap with cell-type specific interactions are reported in (Supplementary file 2).

**Figure 3.**
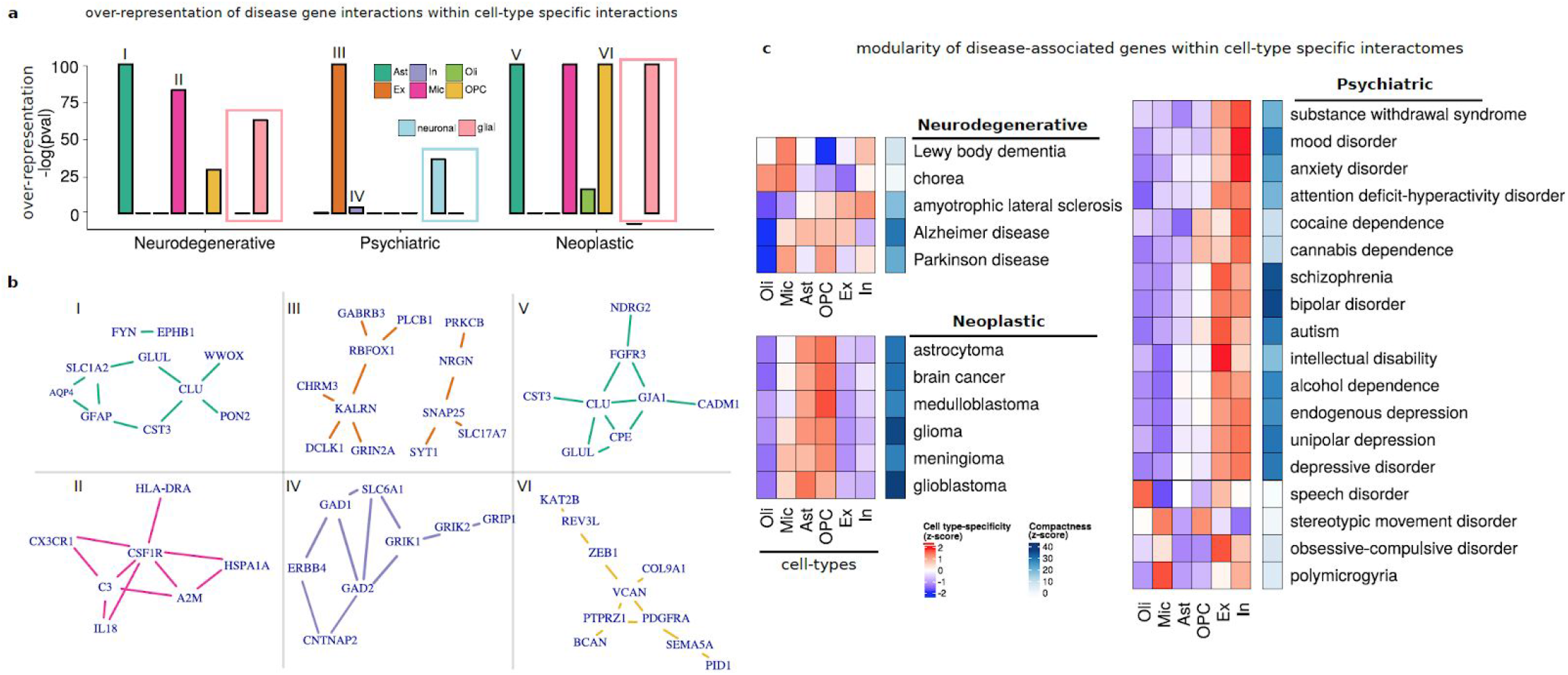
Brain-associated disorders converge to cell-type specific modular perturbations. **a**, over-representation scores for gene interactions between disease-associated genes within cell-type specific interactions are shown for neurodegenerative, psychiatric, and neoplastic disorder groups. Cell-type (Ast, In, Oli, Ex, Mic, OPC) and cell-type group (neuronal, glial) over-representation analysis are considered. Neuronal (blue) or glial (pink) over-representation is highlighted with a rectangle. Roman numbers link cell-type and disease interaction sets with corresponding networks in b. **b**, top 10 strongest interactions for to associated interaction sets considering in a are shows as networks. **c**, network modularity scores for genes associated with individual diseases across cell-types are shown. Modularity is estimated by a compactness score that measures the deviation of module size (average measure of inter-gene network distance) relative to random expectation. Relative (z-scaled) modularity across cell-types is shown in blue-red scale. Maximal compactness for a disease is shown in blue scale.

Thus, our results suggest that genes known to be associated with different disease classes of brain disorders preferentially influence specific cell-types. Interestingly, this approach unbiasedly recovers the increasing realization of a dominant role of inflammatory glial cells (microglia and astrocytes) in neurodegeneration, the known role of neuronal cells in psychiatric disorders, and the expected association of proliferative glial cells (OPC and astrocytes) with brain tumors. In the latter case, our results also suggest an important role of inflammation in neoplasia, as evidenced by the additional overrepresentation of gene interactions occurring in microglia cells.

### Network analysis reveals cell-type-specificity of brain-associated disorders

We next analyzed the modular connectivity of disease genes. We tested whether genes associated with individual brain diseases present non-trivial patterns of interaction across the cell-type specific interactomes; this time considering all genes and interactions in each cell-type network. We followed the rationale of network medicine, which seeks to understand the seemingly independent perturbations commonly associated with a disease in terms of their local and global patterns of connectivity in the interactome (Loscalzo 2017). One way to operationalize such view is through the characterization of disease modules that emerge as dense and tightly compact neighborhoods that topologically localize disease genes within the network (Menche et al. 2015).

Using the set of curated gene-disease associations, we tested for each pair of disease/cell-type network whether disease-associated genes tend to localize in neighborhoods, forming cohesive modules. To quantify *compactness* we used an empirical measure of module size (Loscalzo 2017), defined as the average of the distribution of network distances between pairs of disease genes. To account for the multiplicity of paths between genes, we computed diffusion-based distances (see Methods). Finally, to statistically test whether the observed module size deviates from random expectation, we estimated a corresponding null module size distribution by generating an ensemble of random networks preserving topology while shuffling interaction strengths, and we measured deviation from expectation using a z-score. Overall, we found that disease genes do localize in the interactomes, and that the degree of modularity of disease perturbations varies across diseases (Figure 3c). Across disease groups, neoplastic disorders show the strongest modularity, with glioblastoma genes being the most compacted. Neurodegenerative disorders showed the least modularity, with the exception of Alzheimer’s disease. Neuropsychiatric disorders, on the other hand, showed strong compactness, with SCZ and BPD having the strongest network localization.

Interestingly, we observed highly variable modularity across cell-type specific networks for each disorder, which again points to a pattern of preferential cell-type specificity for the different diseases (Figure 3c). Diseases within the same group show similar patterns of cell-type specificity that are distinct from those observed in the other groups. This observation is consistent with the idea of an association between cell-type convergence patterns of disease perturbations and observed symptomatology, as disorders within the groups more often share patterns and comorbidity than diseases across groups. All neoplastic diseases tested show modular localization in the astrocyte and OPC networks. These two cell-types have proliferative potential and have been discussed in the context of the cell of origin for malignant gliomas (Zong, Parada, and Baker 2015). However, evidence from the specific origin of the tumors in vivo has been proved challenging, given the overlap in gene signatures of astrocytes and neuron progenitor cells. Although we only considered transcriptional states from the adult human brain, SCINET robustly captures the stemness property of the interactomes of astrocytes and OPCs and its association with brain carcinogenesis. Psychiatric disease genes showed strong preferential modular perturbations in neuronal networks, with the exception of disorders of stereotypic movement and speech, which show the strongest compactness on oligodendrocytes. We interpret this observation to be consistent with the myelinating role of oligodendrocytes, as these two disorders involve motor deficiency, which is severely affected in cases of pathophysiology affecting myelin. Indeed, white matter abnormality in the form of delayed or absent myelination has been associated with childhood apraxia of speech (Liégeois and Morgan 2012). Contrasting neuron types, genes associated with mood and depression disorders seem to perturb more strongly the inhibitory neuron network. Different lines of evidence associate deficits in inhibitory neurotransmission to major depressive disorder, such as the reduction of GABA A receptor-mediated cortical inhibition in both late-life and adult depressed subjects relative to controls (Levinson et al. 2010, Lissemore et al. 2018). Finally, unlike the glial/neuronal polarised pattern observed for neoplastic and psychiatric diseases, neurodegenerative disorders showed a more heterogeneous convergence pattern, involving glial and neuronal cell-types in different diseases. This observation might point to the relevance of glial-neuronal interactions, and the conditionality of the protective or damaging effect of glial cell states (De Strooper and Karran 2016). Interestingly, we found that genes associated with Parkinson’s disease converge in both OPCs and neuron networks, consistent with deficiencies on remyelination having an effect on the proper firing of motor neurons.

Overall, our results suggest that the contextualization of disease associated genes within the topologies of cell-type specific networks inferred by SCINET uncovers nontrivial, systems-level interaction patterns among disease genes that provide information on the phenotypic manifestation of the disorder. The consideration of connectivity patterns involving genes which have not yet being associated with a specific disease enables the identification of modular converging perturbations that preferentially occur in specific cell-types.

### Cell-type specific prioritization of Schizophrenia and Bipolar Disorder genes uncovers differential disease components

Even when the dominant pattern of disease gene modular convergence is cell-type specific, we reasoned that a set of disease genes might propagate information differently across the different cell-type networks, a property that could be exploited to potentially find subgroups of these genes with cell-type specific influence. This idea is of especial importance for brain related disorders, which are phenotypically highly heterogeneous, and which likely involve the interaction of multiple causal factors at the cellular level. To explore this idea, we considered the genes associated with both SCZ and BPD (SCZ-BPD genes) as seeds for cell-type specific network propagation. Using a modified symmetric random-walk algorithm (see Methods), we ranked all the genes in each cell-type network based on their proximity to the seed genes -- a practice commonly referred to as gene prioritization (Guala and Sonnhammer 2017). Given the differential interaction strengths across cell-type networks, the resulting gene ranking varies across cell-types. In line with the highly heterogeneous phenotypic manifestations of the disorders, we hypothesized that prioritization could uncover different subsets of SCZ-BPD genes that are topologically influential in different cell-types. In order to directly explore this, we extracted from the reference interactome the subnetwork defined only by the shared genes and their corresponding interactions (SCZ-BPD subnetwork). Subsequently, we projected to this network the prioritized cell-type specific gene rankings and the interaction strengths, resulting in 6 cell-type specific SCZ-BPD subnetworks. We found that, indeed, different subsets of the genes rank high and localize in the different cell-types, suggesting a differential topological influence across cell-types (Figure 4a).

**Figure 4.**
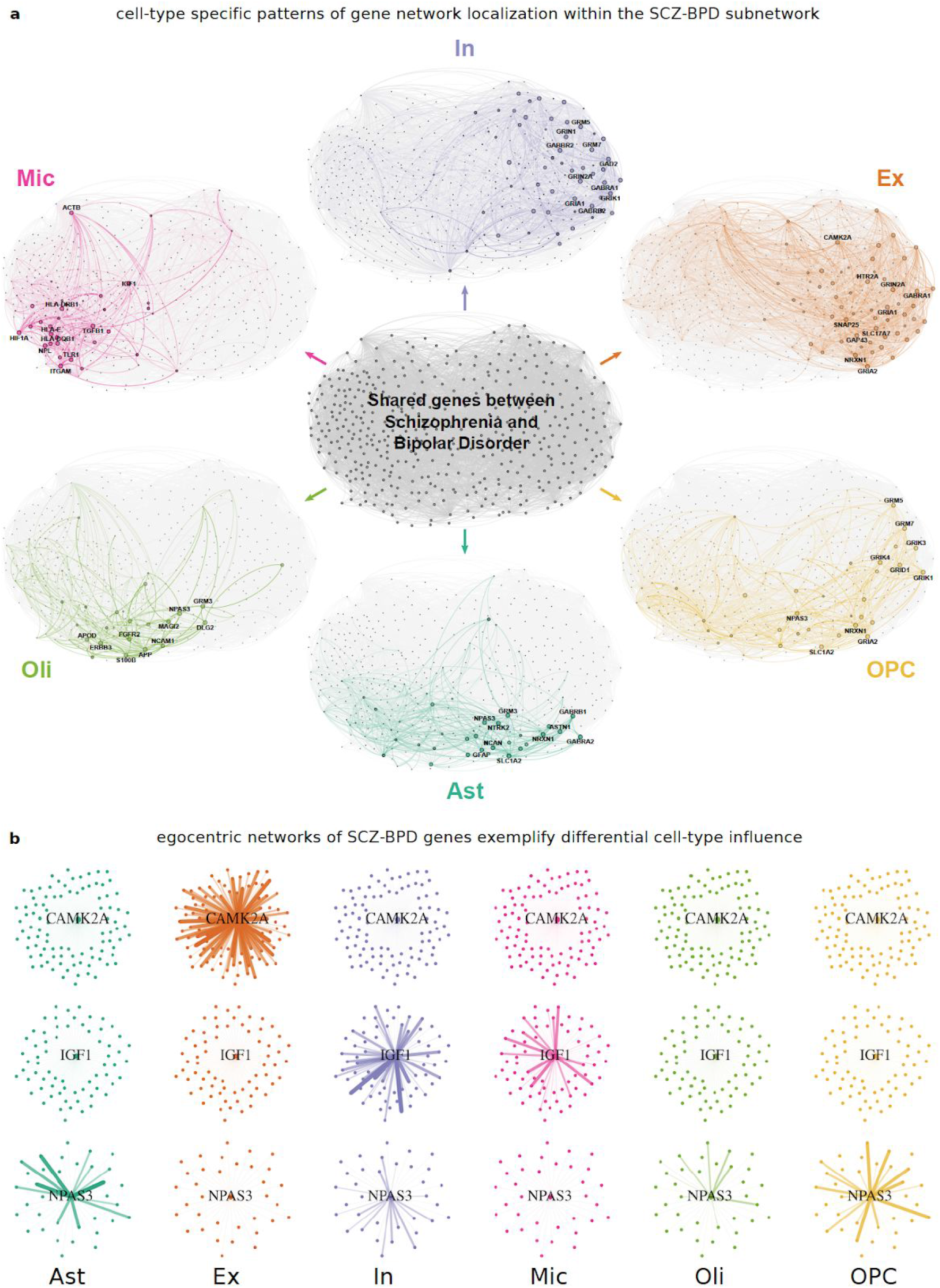
Cell-type specific prioritization of genes associated with both SCZ and BPD (SCZ-BPD genes). **a**, SCZ-BPD interactome subnetwork constructed by extracting SCZ-BPD genes and their mutual interactions from the human reference interactome. After network propagation analysis over each cell-type specific interactome, the resulting gene prioritization scores and SCINET inferred interaction strengths are projected to the SCZ-BPD subnetwork, independently for each cell-type. Top 10 genes based on prioritization score are highlighted. Node size and color transparency are proportional to scores. **b**, egocentric networks for selected genes. Each egonet shows a selected gene (center) surrounded with its direct interacting partners. Edge width is proportional to interaction strength. Genes display strong interactions only in a small subset of cell-types.

Overall, genes involved in the pathways with well-established molecular roles in psychiatric neurobiology consistently ranked among the top prioritized genes in associated cell-types. For example, we found that glutamate signaling genes (GRIN1, GRIN2A, and GRIN2B) and AMPA receptors (GRIA1 and GRIA2), are highly influential in the neuronal networks. For glial cells, we found that NPAS3, which plays a regulatory role in neurogenesis (Sha et al. 2012; Yang et al. 2016), influences the networks of cell types with proliferative capacity (OPC and astrocyte). In microglia we found Hypoxia-Inducible Factor (HIF)-1A, a gene that responds to a decrease in cellular oxygen availability, and which has been shown to mediate microglial activation affecting neuron function (Lu et al. 2006). Finally, neurexin 1 (NRXN1), a gene whose genetic perturbations are strongly associated with psychiatric disorders (Duong et al. 2012) and which is involved in synaptogenesis and synapse maintenance (Michaelson et al. 2017) is influential in both neuronal (excitatory) and glial (astrocyte and OPC) networks. These observations show that the topologically influence of a disease gene estimated through its patterns of cell-type specific connectivity is able to point to the cellular context of its effect. The discovery of multiplicity of influence across cell-types also suggest that comparative propagation analysis might recover the effect of genes in glial-neuronal interactions. To facilitate the future study of unknown but potentially relevant SCZ-BPD genes, we report cell-type specific prioritization results in Supplementary Table 4.

To more directly explore the differential influence of SCZ-BPD genes across cell-types, we further defined gene-centered egocentric networks (egonets) (Figure 4b). These networks are commonly used in social network analysis (Karsai, Perra, and Vespignani 2014) and consists of a center node and its direct interacting partners, which provides an intuitive way to directly visualize the dynamics of gene influence. We found two dominant patterns of interaction: (i) genes having an exclusive influence in one cell type, and (ii) genes simultaneously influencing multiple cell-types (Figure 4b). For example, we can clearly see that CAMK2A, a key kinase in calcium signaling, exclusively influences excitatory neurons; while NPAS3, a brain-enriched transcription factor, mainly has strong interactions in OPC and astrocyte networks. On the other hand, IGF1, a hub in the microglia-specific network, exhibits a strong influence in both microglia and inhibitory-neuron networks. Gene set enrichment analysis of the IGF1 neighbors in microglia and inhibitory neurons, independently uncovers enriched processes consistent with a systemic, inter-cellular role. In the microglia network, IGF1 interacts with genes involved in the regulation of neuron death (p-val 4.71e-07, hypergeometric test) and response to hypoxia (1.70e-05), including major regulators linking metabolism and protein homeostasis such as BDNF and BCL2. In addition, IGF1 also interacts with genes involved in the circadian rhythm (2.96e-08), a systemic process known to feedback with the levels of BDNF and its neuroprotective function (Marosi and Mattson 2014).

Thus, while disease genes tend to predominantly form active modules in specific cell-types, the cell-type specific interaction strengths inferred by SCINET enable the identification of specific subgroups of genes with preferential influence on different cell-types. Gene prioritization of the same set of genes across multiple condition-specific SCINET networks, facilitates the contextualization and discovery of unknown but potentially relevant target genes.

## Discussion

We introduced SCINET, a novel computational framework that enables the reconstruction of cell-type specific interactomes by leveraging single-cell transcriptomic data. By inferring and quantifying cell-type specific gene interaction strengths, SCINET provides a cellular context to interpret molecular pathways and functional modules. SCINET can be used to contextualize disease associated genes and their quantifiable influence in different cell-types or conditions, to study potential mediators of functional interactions between cell types, or to compare interaction strength distributions and assess the dynamics of interaction usage across developmental or pathological conditions. SCINET is of general applicability and can be used to analyze any combination of cell conditions and reference (physical and/or functional) interactomes. We envision SCINET as a simple to use computational tool to aid in the design and interpretation of cell-type resolution experiments and to uncover context-specific convergence of heterogeneous and seemingly independent genetic perturbations.

## Materials and methods

### Data sources

#### Human interactome

The Parsimonious Composite Network (PCNet) (Huang et al. 2018) was used as reference human interactome. This network was constructed by combining 21 heterogeneous network resources, including STRING, ConsensusPathDB, and GIANT, among others. PCNet retains edges that are supported by at least two independent sources, which leads to a high-sensitivity and high-specificity network that outperforms any individual network with respect to predicting disease-associated genes (Huang et al. 2018). PCNet was used throughout our paper, and we refer to it as the “global human interactome”.

#### Single-cell atlas of human prefrontal cortex

A recently published dataset of single nuclei RNA sequencing from the human frontal cortex (9 male, 4 female individuals) was used as reference transcriptome. After quality control, this dataset contains a total of 10,319 single cells (4,164 and 6,155 cells from BA6 and BA10 regions, respectively). Six major cell-types of the brain were considered: astrocytes, excitatory/inhibitory neurons, microglia, oligodendrocytes, and oligodendrocyte progenitor cells (OPCs). All subtypes of excitatory and inhibitory neurons were independently combined to create cell-type annotation for these two classes.

#### Cell-type specific gene markers

Cell-type-specific marker genes as defined by Lake et al. were used. Only genes with at least 1-log-fold-ratio difference when cells of a given cell-type are compared against the rest of cells were considered. This resulted in marker genes for astrocytes (79), excitatory neurons (162), inhibitory neurons (304), microglia (45), oligodendrocytes (103), and OPCs (52).

#### Compendium of genes associated with various brain disorders

Disease-associated genes were collected from the DisGeNET database (Piñero et al. 2017), which aggregates data from GWAS catalogues, animal models, and the scientific literature; preserving the evidence type supporting each disease-gene association. All disorders were mapped to the Monarch Disease Ontology (MonDO) (Bello et al. 2018) and brain-associated disorders corresponding to (i) neurodegenerative, (ii) neuropsychiatric, and (iii) brain cancers were selected. To this end, first diseases that are associated with nervous system (annotated with “nervous system disorder (MONDO_0005071)” term) were selected. Then selected disorders were intersected with disorders annotated with “neurodegenerative disease” (MONDO:0005559), “psychiatric disorder” (MONDO:0002025), and “neoplastic disease or syndrome” (MONDO:0023370) terms. The final dataset contains 5 neurodegenerative disorders, 18 neuropsychiatric disorders, and 6 types of brain cancers.

### Interpolation and smoothing of expression profiles

Archetypal analysis for Cell Type identificatION (ACTION) (Mohammadi et al. 2018) characterizes the transcriptional landscape of single cells using an optimal set of archetypal states. At the core of this method there is an optimization framework to identify characteristic landmark cells that can be used to optimally represent the rest of cells. Formally, given an expression matrix **S** ∈ ℝ^*genes×cells*^ ACTION identifies a set of *archetypal states* that optimally represent the rest of cells. Each of these archetypal states are a convex (sparse) combination of cells in the data. In this regard, ACTION reduces the noise from dropouts by local averaging and smoothing of cells, while preserving the subtle state differences by enforcing a sparsity constraint on the number of cells that are being averaged. The original version of the ACTION method was based on a kernel-based formulation. To deal with the rapidly growing scale of the single-cell profiles, we implicitly reduce this kernel (of dimension cell × cell) to another subspace, **S**_*r*_ ∈ ℝ^*D×cells*^ such that the dot-product of columns in the reduced subspace recapitulate the kernel matrix. To this end, we compute the SVD decomposition of the orthogonalized expression profile (based on the ACTION method) as 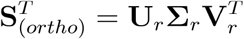. Then, we compute the reduced expression profile as 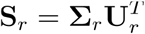, in which every row represents a *metagene* and each column represents a cell. Then the following optimization problem is solved to find a set of landmark cells using *convex non-negative matrix factorization (convNMF)*:

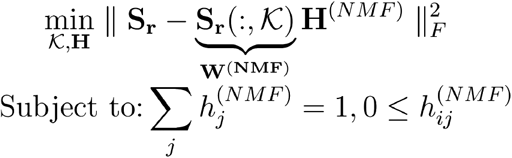

where 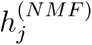 is the *j^th^* column of the *H*^(*NMF*)^ matrix. Using these initial landmark cells encoded in the matrix *W*^(*NMF*)^ we then initialize a round of *archetypal analysis (AA)* to smooth and denoise them and compute the final archetypal states. Mathematically, we solve the following optimization problem:

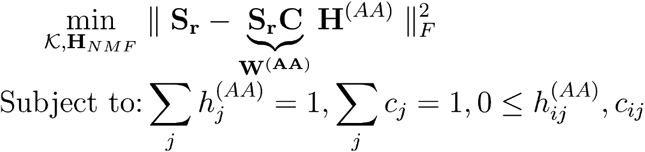

By initializing matrix **W**^(*AA*)^ = *W*^(*NMF*)^ this formulation enables the smoothing of the profile of archetypal cells based on a small number of close-by cells, as sparsity is enforced due to norm-1 constraint on the columns of C.

Another modification to the original formulation is that in ACTION the total number of archetypes 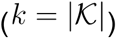 was fixed. However, different cell-types/states are best captured at different levels of resolutions, and thus different values of *k* could be optimal for identifying different transcriptional patterns. To address this issue, increasing values of *k* are allowed by running ACTION at multiple resolutions and then the set of all archetypes across resolutions are combined. The resulting set of archetypes is refer to as the *multi-level archetypal set* (**W**^(*ML*)^). Subsequently, the matrix **H**^(*ML*)^ is recomputed by regressing over the **W**^(*ML*)^ matrix. Given the resulting multi-level archetypal profiles, which are of dimension metagene × aggregate archetypes, a reverse projection onto the state space of genes is computed using matrix **V**_*r*_. Then, for each cell, the matrix **H**^(*ML*)^ is used to interpolate its corresponding expression profile using the archetypes profile. Putting it all together, the described algorithms is implemented in matrix operations as follows:

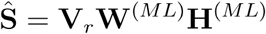

in which **H**^(*ML*)^ is computed using the multi-level archetypal set, **W**^(*ML*)^ This approach is fundamentally different from common gene imputation methods in that signature genes, which distinguish cell-types, will have high-values in the interpolated profiles, even when their absolute expression value is small. Furthermore, for larger datasets, given the flexibility of matrix computations, it is possible to interpolate values for only a subset of genes and/or cells of interest -- i.e., by extracting and using in the operation only the corresponding rows/columns.

### Adjusting interpolated profiles for the skewed distribution of gene expressions values

The feasibility of an interaction in a given context is assessed by comparing and combining the expression value of each pair of interacting genes. However, different genes have radically different expression distributions. Moreover, the baseline expression, corresponding to the mean of these distributions, is not comparable, since some genes might be functional at much lower doses than others. To address this issue and to put expression measurements on the same scale, the *rank-based inverse Normal transformation* technique was used. Given the interpolated expression profile Ŝ with *m* genes and *n* cells, two independent factors are computed, one for the expression of each gene across cells (row factor) and one for the mean expression of all genes in a given subpopulation of cells (column factor). In the former, given a gene *i*, its expression profile is sorted across all cells and a rank *r_ij_* is assigned to each interpolated expression profile *Ŝ_ij_*, which is then normalize by the total number of cells, 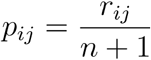. The row-factor matrix, **F**^(*r*)^ is then computed by projecting the normalized ranks onto the standard Normal distribution: 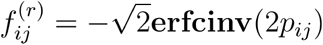, where **erfcinv** is the inverse of the complementary error function, **erfc**, defined as:

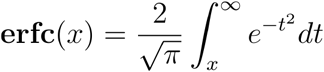

Similarly, a column factor for all genes is defined. Given a subset of cells, 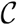, representing the cell-type of interest, columns of interpolated expression profile are averaged within the subspace of 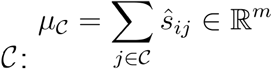. Then, a column factor vector 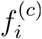 is defined by transforming 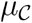 in a fashion similar to the row factor. Finally, using the two components, a transformed, interpolated expression profile matrix, **T**, is defined in which

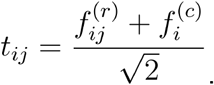

### Assessing co-expression dependencies between pairs of interacting proteins

Our working assumption is that an interaction can only happen if both endpoints of the interaction are expressed at a high enough levels. Such notion can be formalized by requiring the minimum expression of two interacting proteins to be “large enough”. Given that the expression value of each gene is transformed to follow a standard normal distribution, under null assumption of independence between the expression value of two genes, the right tail of the min operator can be computed using:

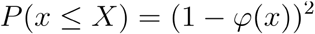

where *x* = *min*(*t_ij_, t_i’j_*), in which *t_ij_* and *t_i’j_* are the transformed expression values of gene products corresponding to vertices *i* and *i’* incident on an interaction *i* − *i*’ in cell *j*, and *φ* is the cumulative density function (CDF) of the standard normal distribution.

This definition allows the computation of a *p*-value for each interaction in a given cell. To account for noise and different sample-size across cell-types a technique similar to ensemble learning was adopted by selecting *k* cells at random, computing their individual interaction *p*-values, and combining these *p*-values into an aggregate metap-value using the Fisher’s combination method. Specifically, denote by 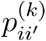 the *p*-value of interaction *i* − *i*’ occurring in the *k^th^* sampled cell, then an aggregated p-value statistic is computed by:

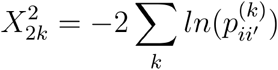

When the null hypothesis of each individual test is true and the tests are independent, 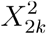 follows a *χ*^2^ distribution with 2*k* degrees of freedom, which can be used to compute the meta p-value associated with all tests. This subsampling scheme balances the total number of cells in the given population.

Finally, we note that by repeated application of the resampling method, an empirical distribution over each gene interaction that described its dynamic characteristics across cells can be estimated. The first moment (mean) of this distribution is used for analyses reported in the paper.

### Topological specificity

A measurement is introduced to decouple the global, context-agnostic connectivity of a gene as provided by its number of interactions in the global interactome, from the cell-type-specific strength of interactions incident to the gene in cell-type-specific network. First, the hubness of each gene in a given cell-type-specific network is computed as the total strength of its local neighbors, represented by *w*^(*celltype*)^(*i*) for each protein *i*. Second, a random model is used to estimate the deviation of this observed hubness relative to random expectation. The random model consists of an ensemble of networks in which the underlying topology of the global interactome is preserved while the cell-type-specific edge weights are reshuffled uniformly at random. For each random network, the strength of interactions is recomputed, resulting in a distribution of gene neighborhood strengths for each gene. Using the mean and standard deviation of each distribution 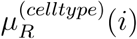 and 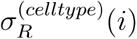, respectively; the topological-specificity of each gene in a given network is define as:

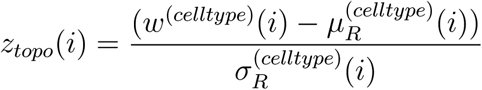

### Compactness and prioritization of disease-associated genes

Random-walk methods are effective techniques to establish network-based relationships among disease-associated genes (Köhler et al. 2008). Here, a symmetric version of the random-walk with restart process is used to prioritize disease genes (Vanunu et al. 2010). More specifically, given a cell-type-specific network represented using its adjacency matrix, **A**, the following stochastic, symmetric transition matrix is defined:

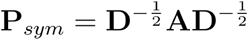

where 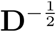 is a diagonal matrix with the strength of each node (column sums in **A**) as diagonal elements. Using this transition matrix, the stationary distribution of the random-walk process is defined in closed form as:

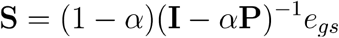

where *e_g_s* is a stochastic vector of restart probabilities and *α* a parameter that adjusts the depth of the random-walk process ((1 − *α*) is the probability of starting a new random-walk from one of the seed nodes in *e_gs_*). To compute the *α* parameter, an approach similar to the one proposed in Huang *et al. (Huang et al. 2018)* was used. Briefly, the optimal choice of *α* is set using the following linear model against the log10-adjusted number of interactions in the network (*nE*):

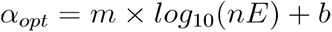

where *m* = −0.03 and *b* = 0.75.

Finally, we define the restart probabilities encoded in vector *e_gs_* based on the topological-specificity of the corresponding proteins as follows:

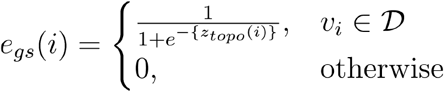

where 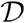 is the disease geneset of interest. The resulting vector was normalized by its sum to construct a stochastic vector to be incorporated in the random-walk.

### Availability

C++ implementation of SCINET with R/Matlab interfaces is freely available at: https://github.com/shmohammadi86/SCINET.

## Supplementary Figures

**Supplementary Figure 1.**
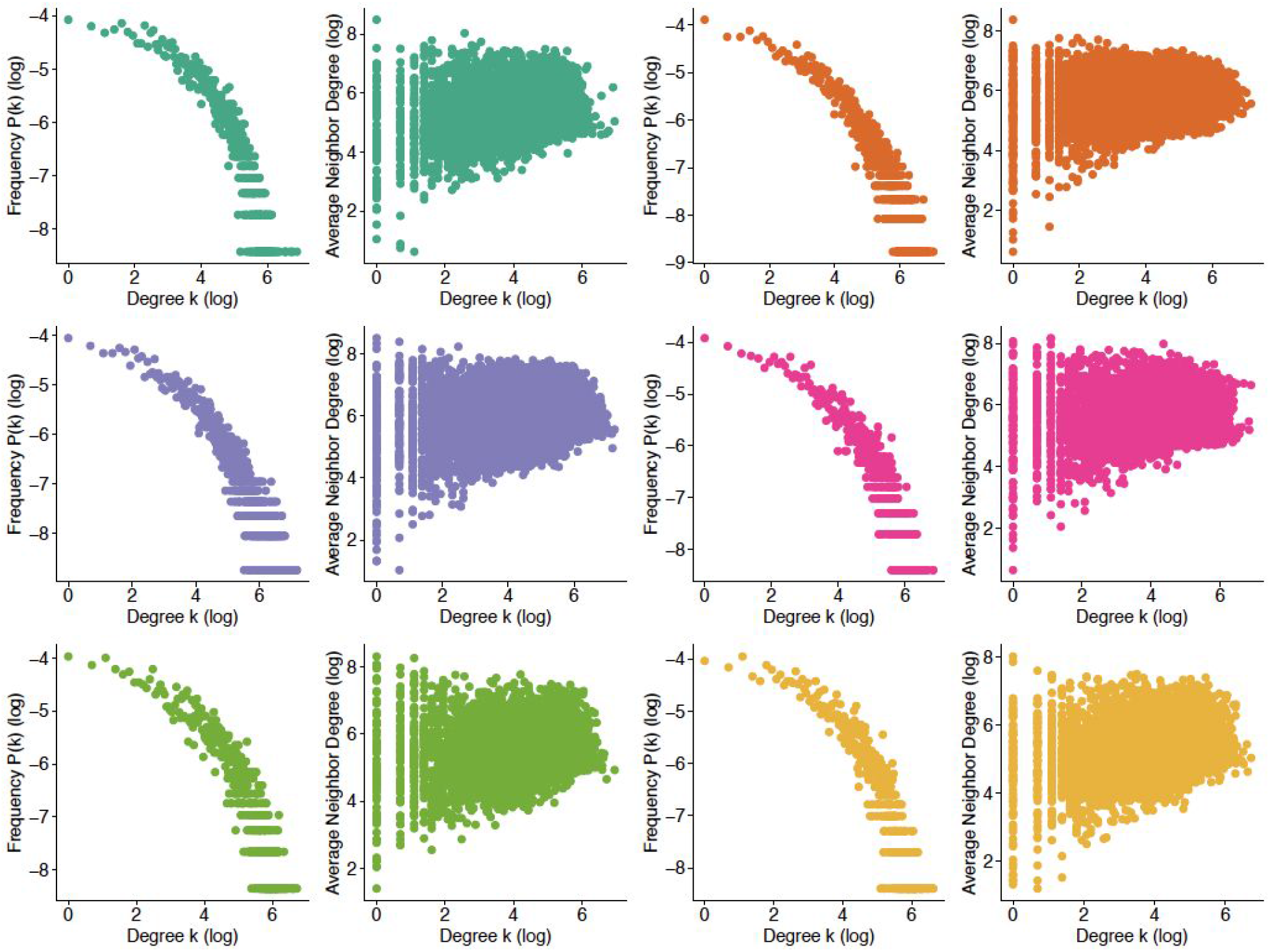
Gene connectivity and partner preference distributions across cell-type specific interactomes. Degree (total number of interactions of a given gene) distributions are displayed in log scale (left). Patterns of gene neighbors’ connectivity as a function of a gene’s degree are displayed in log scale (right). Overall, connected genes show a slight preference to interact with genes.

**Supplementary Figure 2.**
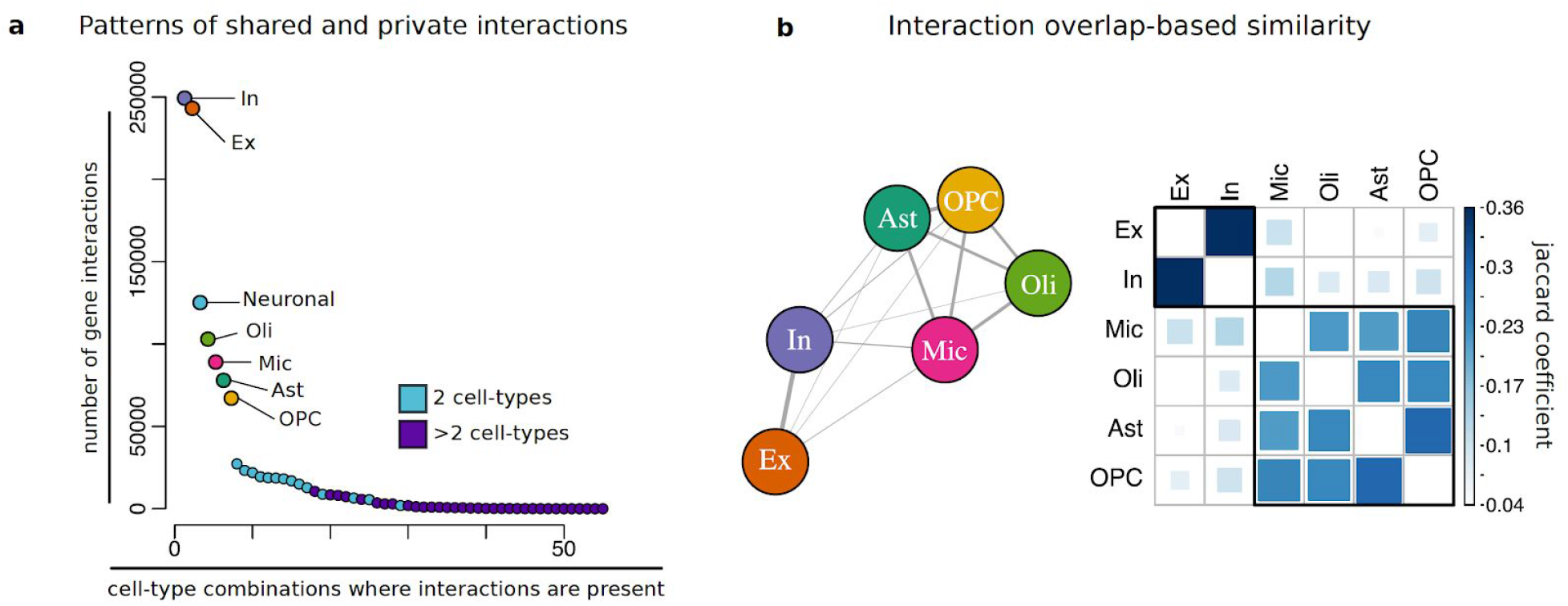
Patterns of cell-type shared and private interactions. **a**, number of gene interactions that are observed in only one cell-type (cell-type colors), only two different cell-types (light blue), or more than two different cell-types (purple). Cell-type combinations refers to any combination of one of more cell-types in which a gene interactions occurs. **b**, Network representation linking cell-types based on the degree of overlap in shared gene interactions (left). Cell-types that share more interactions appear positioned close together. Edge weight is proportional to the degree of overlap. Degree of overlap in interactions is quantified using the pairwise jaccard coefficient. Corresponding Jaccard coefficients are shown in matrix form (right).

**Supplementary Figure 3.**
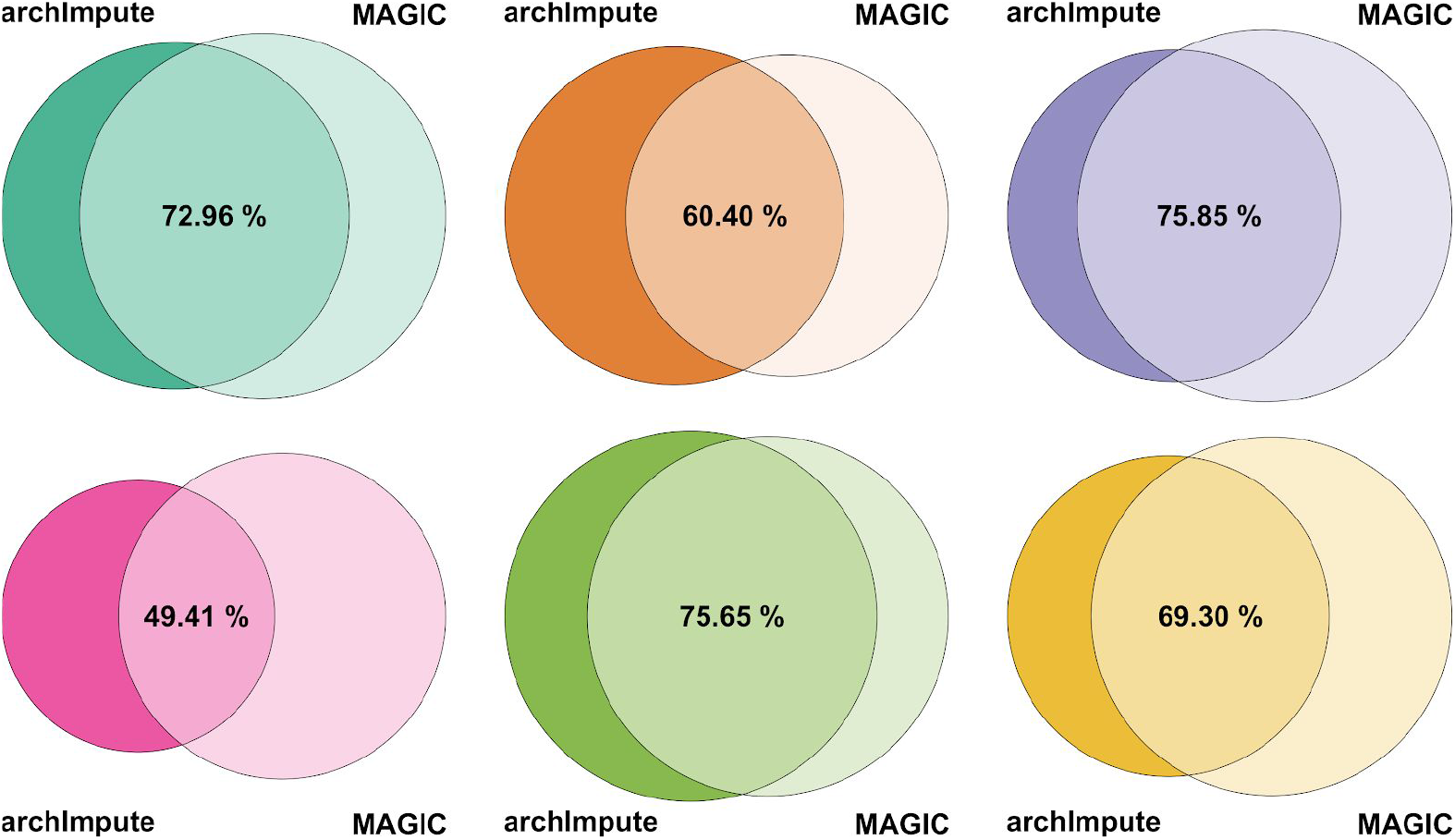
Robustness of SCINET to choice of imputation method. Each panel shows the overlap of identified edges, after thresholding p-values (0.05), across different cell types. Except in microglia, the majority of predicted edges overlap with each other, with the strongest overlap observed in inhibitory neuron and oligodendrocyte-specific networks. Expression values for genes were imputed, independently, using either ACTION-based archetype imputation (archImpute, see methods) or MAGIC method. Imputed profiles were used to infer cell-type specificity of edges.

**Supplementary Figure 4.**
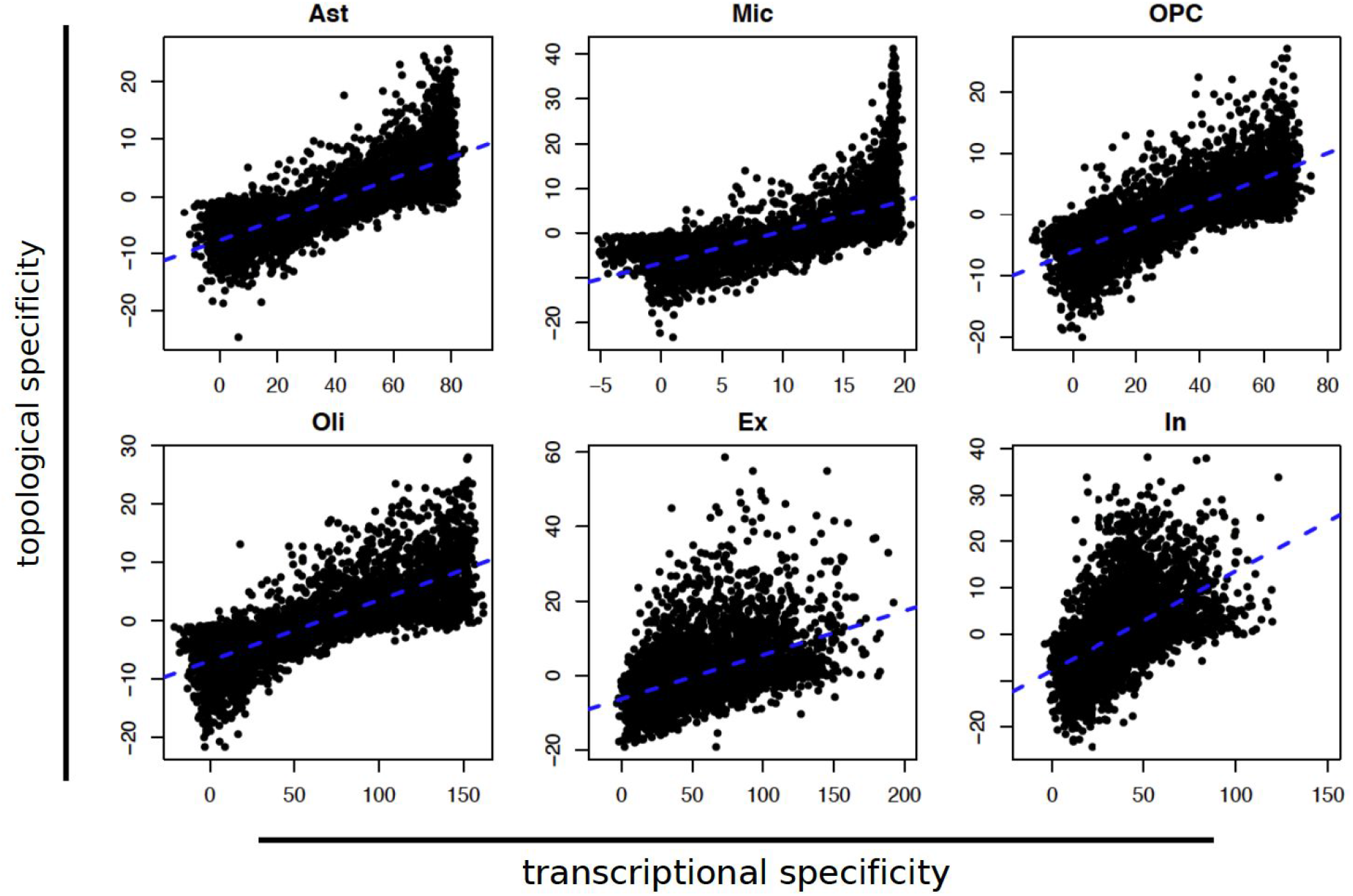
Global consistency of topological and transcriptional specificity. Plots show corresponding topological and transcriptional specificity values across genes of each cell-type specific network. Blue line represents the best linear regression fit.

